# *In Vitro* Anti-cancer potential of Aloe emodin against Oral Squamous Cell Carcinoma

**DOI:** 10.1101/2024.04.12.589239

**Authors:** Jesse Joel T, Jagadish Kumar Suluvoy, Harish babu Kolla

**Affiliations:** Department of Biotechnology, School of Agriculture and Biosciences, Karunya Institute of Technology and Sciences (Deemed to be University), Coimbatore, Tamil Nadu, India; Department of Biotechnology, Vignan’s Foundation for Science Technology and Research, Guntur, Andhra Pradesh 522213, India

**Keywords:** Oral cancer, Aloe emodin, *in vitro*, anti-cancer, KB 3-1 cell line

## Abstract

Oral squamous cell carcinoma is caused due to smoking, chewing of tobacco, high exposure of UV radiation and infectious agents such as human papilloma virus (HPV). Epigenetic dysregulation and uncontrolled cell growth are main factors that drives towards tumor development. Risk factors such as smoking, tobacco and viral agent’s up regulates its activity and leads to the cancer. Although chemotherapy, surgery and radiation were preferred for treating cancer but patients, feel extreme psychological stress and depression. Natural products are been investigated for their numerous diseases and in that they can be treated against cancer without side effects. Aloe emodin is one such natural molecule available, which possess various pharmacological activities for human beneficial. In this current study, we investigated the anti-cancer activity of aloe emodin *in vitro*. The KB 3-1 cell line were treated with different concentrations of Aloe emodin and its anti-cancer potential was determined through DNA fragmentation and cell viability assays.

## Introduction

Oral cancer is highly prevalent in India and other developing countries. This incidence is due to the most common habits of smoking and chewing tobacco. Oral cancer remains a challenge to patients because of its pathogenicity and various adverse reactions being faced by patients during the treatment. This condition is responsible for impairment of day-to-day activities viz. eating, swallowing, breathing, and speaking.

Squamous cell carcinoma (SCC) accounts for nearly 90% of all oral cancer cases. It may affect any anatomical site in the mouth, but most commonly the tongue and the floor of the mouth. Oral SCC pathogenicity increases with the prolonged exposure to risk factors, and aging that adds a dimension for age-related mutagenic and epigenetic changes. Besides these risk factors, infectious agents such as human papilloma virus is an important agent that causes oral cancer [1] (Herrero et al., 2003).

Chemotherapy, radiation, and surgery are generally used to manage squamous cell carcinoma, but these medications are not the complete cure. Although various chemotherapeutic alternatives were available for treating cancer, their adverse side effects have become disadvantages. Patients suffer from severe fatigue, anxiety and depression during the treatment and can persist for long periods of time [2] (Goswami et al., 2019).

Currently, the interest towards cancer therapy is shifting to natural compounds and phytochemicals to fight against various diseases. Natural products mainly from plant source investigated for their extensive biological activities with health benefits. Emodin was one such molecule with anti-cancer, anti-diabetic and anti-microbial activities. Emodin was first isolated from a Chinese medicinal plant *Rheum palmatum, Cassia occidentalis* and *Polygonum multiflorum*. The overall molecular mechanism of emodin includes arrest of cell cycle, anti-apoptosis, anti-proliferative on cancer cells [3-5] (Yu et al., 2013; Tabolacci et al., 2010; Vath et al 2002). Along with this, Emodin has anti-inflammatory, antiviral, antibacterial and anti-parasitic [6] (Dong et al., 2020). Emodin fights against surplus inflammation by selecting and attacking miR139-5-LO and balancing the factor 2 of erythroid and glycogen synthase kinase 3β. This indirectly explains the emodin as the most promising compound to treat oral squamous cell carcinoma [7] (Zhang et al., 2020).

Due to its wide therapeutic medicinal and pharmacological activities, our interest was to study the anti-cancer activity of Aloe emodin, an isomer of emodin through *in vitro* and *in silico* approaches. In the current experimental research cytotoxicity, anti-proliferative, DNA fragmentation investigated in *in vitro* studies an through computational approach, inhibitory actions of aleo emodin, an exudate from aloe and its pharmacological nature is studied.

## Materials and methods

### Cell line and reagents

Human KB 3-1 cell line was procured from National Center for Cell Science (NCCS) Pune, Maharashtra, India. The cells were cultured in DMEM media supplemented with 10 % FBS and maintained in 5 % CO2 incubator at 37°C. DMEM media, Aloe emodin, all the chemicals and reagents used in the study were analytical grade and procured from Sigma Aldrich, India.

### Cell viability assay

The cell viability assay was performed in duplicates with appropriate controls. KB 3-1 cells were treated with Aloe emodin at various concentrations of 100 μg/mL, 125 μg/mL and 150 μ/mL g. Untreated cells were considered as control. Cells were allowed to grow at 37 °C for 72 hrs in a 5 % CO2 incubator. Cell viability was assessed through trypan blue assay by taking an aliquot of cell suspension and centrifuged at 1000 rpm for 5 minutes. Supernatant was carefully discarded, and 5 mL of trypsin added to the flask and set aside for 5 min until the cells detach from the flask. Cells were washed twice with sterile 1X PBS and mixed with 0.4 % of trypan blue in equal proportions and incubated for three minutes at room temperature. Total cells were counted using a hemocytometer. Number of viable cells were calculated with the formulae:

Viable cells % = (total no of viable cells per ml /total no of cells per ml) ^*^100

### DNA Fragmentation

DNA fragmentation in the KB 3-1 cells was studied in duplicates to determine the potential of Aloe emodin in cell death. Different concentrations (100 μg/mL, 125 μg/mL, and 150 μg/mL) of Aloe emodin were added to different wells containing KB 3-1 cells in DMEM media. Untreated cells are considered as control and cells allowed to grow for 72 hrs. After incubation, cells from both the treated and control wells were centrifuged and washed twice with sterile PBS and DNA has been isolated through boil lysis from the cells and the OD values are recorded at 260 nm.

## Results

### Cell viability Assay

Cell viability was assessed based on the principle that dead cells accept trypan blue and the dye passes through the membrane. Percent of viability of KB 3-1 cancer cells was higher in control with an average of 95.75%. Similarly, the inhibition is also very minimal in controls at an average of 4.25 %. The percentage of viable cells decreased with increase in the concentration of Aloe emodin. Minimum viability and maximum inhibition of KB 3-1 cell growth was recorded at 150 μg/mL concentration (Figure 1 and 2).

**Figure 1.**
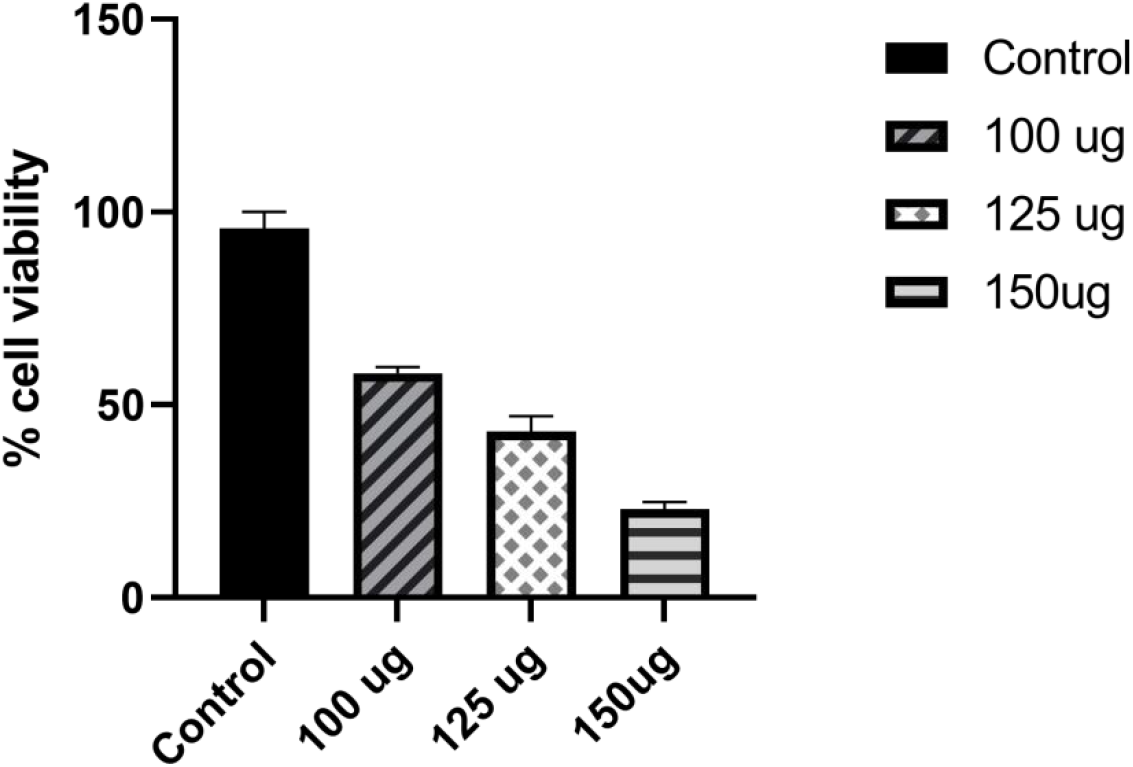
Cell viability in control and KB 3-1 treated cells. Aloe emodin exhibited maximum inhibition at 150 concentration.

**Figure 2.**
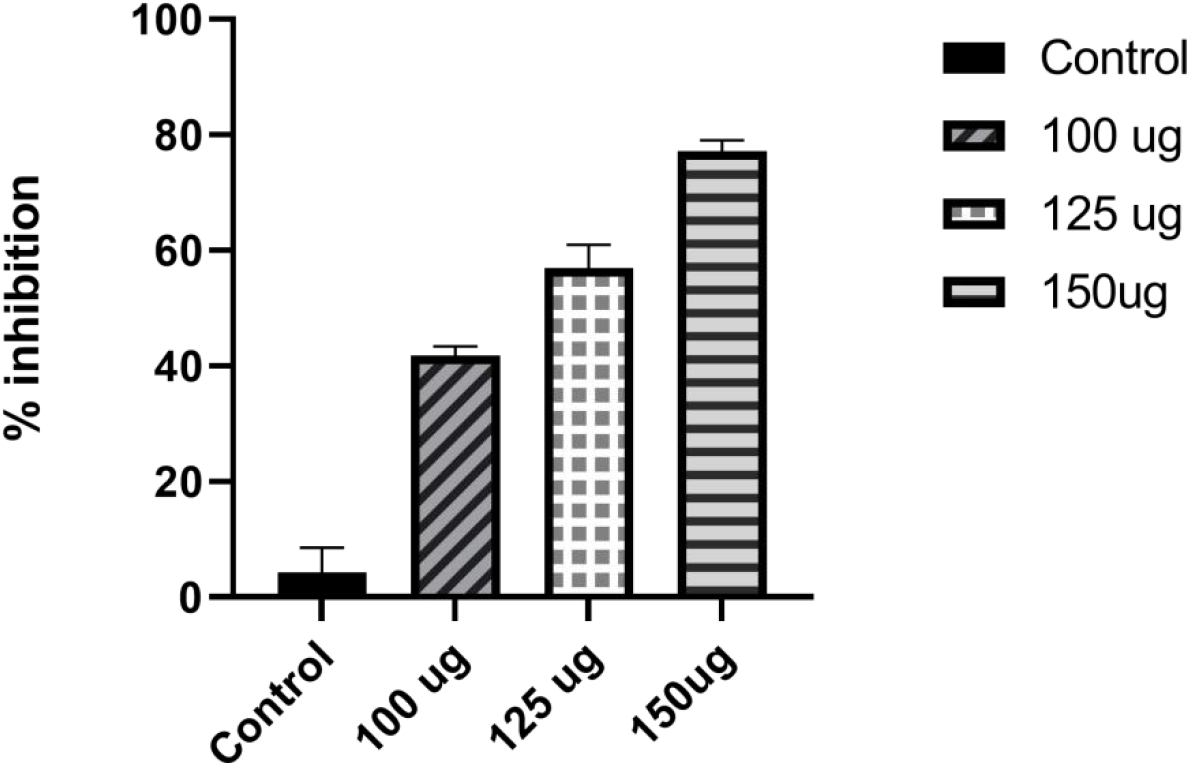
Percentage inhibition of cell growth in control and treated KB 3-1 cells. Maximum inhibition is seen in with the increased concentration of Aloe emodin.

### DNA fragmentation

DNA fragmentation or damage was very crucial during the cell death and apoptosis. Any anti-cancer compound induce its anti-cancer activity by damaging the cell’s DNA and leading to apoptotic death. Extent of DNA fragmentation was studied in Aloe emodin treated KB 3-1 cancer cells. Damaged DNA fragments absorb much UV light than high quality one. Average OD values were found to be high in treated group cells that were treated with different concentrations of Aloe emodin than control (Figure 3). Highest OD value was observed for KB 3-1 cells treated with 150 μg/mL of Aloe emodin (Figure 3).

**Figure 3.**
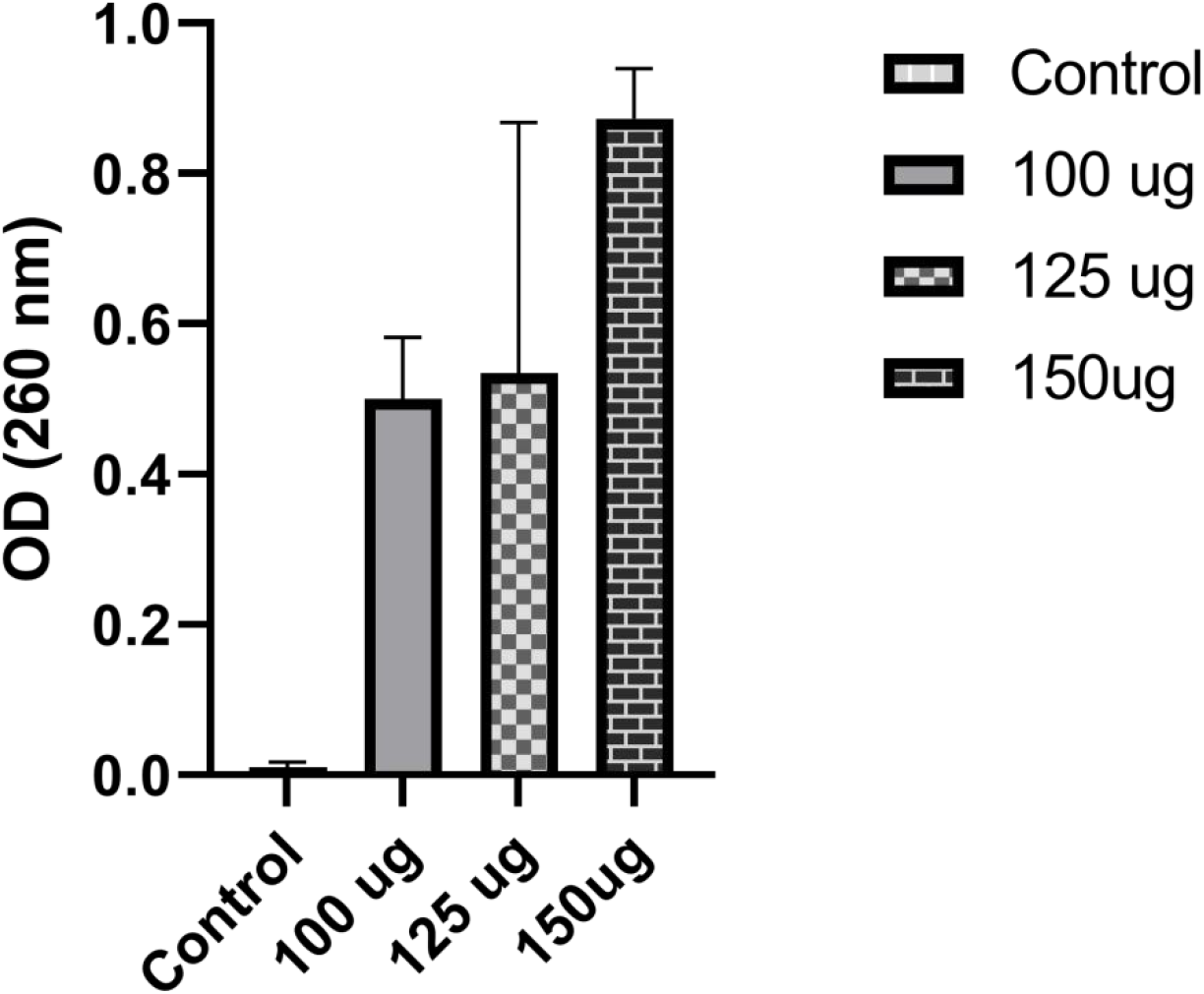
DNA fragmentation in KB 3-1 cells due to Aloe emodin. The increase in the concentration showed the increase in DNA fragmentation.

### Drug likeliness

Drug likeliness for any compound was investigated based on Lipinski’s five rules viz. Molecular weight <500 Da, < 5 H bond donors, < 10 H bond acceptors, LogP<5 and molar reactivity of the molecule should be in the range of 40-130. For any compound to be an ideal drug, it must satisfy all the five Lipinski’s rules i.e., <500 Da in molecular weight, <5 H bond donors and <10 H bond acceptors, molar reactivity in the range of 40-130 and LogP value <5.

Similarly, the drug likeliness properties of Aloe emodin were determined using Lipinski’s rule of five [8] (Lipinski, 2004). Molecular weight of Aloe emodin was 270 Da. Number of H bond donors and acceptors were 3 and 5. LogP and molar reactivity of the molecule were 1.365500 and 69.001381 respectively. Aloe emodin satisfied all the five rules of Lipinki’s and found as an ideal drug molecule.

## Discussion

Cancer therapy with the natural products is gaining importance to cure various cancers because of their low toxicity and side effects as compared to the conventional chemotherapeutics or radiation therapy. Aloe emodin is such a powerful natural compound that is obtained from the Aloe vera and posses various biological activities such as anti-microbial and immunomodulatory products [6].

We studied the *in vitro* anti-cancer potential of Aloe emodin against the oral squamous cell carcinoma on KB 3-1 cell line. Our findings clearly shows the greater decrease in the viability of cancer cells in treatment group as compared to control. Similarly, we have seen the action of Aloe emodin on the KB 3-1 cells through DNA fragmentation assay where the DNA was degraded which is a key mechanism of any anti-cancer drug.

In supporting with the *in vitro* results, drug likeliness properties of Aloe emodin was determined through *in silico* analysis. These results also show that the Aloe emodin as a potential drug candidate by satisfying all the 5 rules of Lipinki’s.

We strongly believe that our initial findings will provide a path for the current research in natural product research to shed a light on Aloe emodin as a potential anti-cancer candidate against the OSCC and beyond. However, further animal studies are needed to be carried out to study the anti-cancer potential of this compound in depth.

## Conclusion

Anti-cancer activity of Aloe emodin was investigated *in vitro* and its drug likeliness properties was studied *in silico*. Aloe emodin exhibited its cytotoxicity and inhibited the growth of KB 3-1 cells. Apoptotic and cell death due to the compound was assessed through DNA damage assay. *In vitro* studies on KB 3-1 cell line has proven that Aloe emodin as a potential anti-cancer molecule. Drug likeliness properties and affinity of the compound were studied through *in silico* methods. Aloe emodin exerted its anti-cancer activity on KB 3-1 cell line in *in vitro* studies and identified as a potential drug molecule owing for its drug likeliness.

## Disclosure statement

No potential conflict of interest was reported by the authors.

## Funding

Not applicable.

## Acknowledgement

The authors thank management of Karunya University and Vignan’s Foundation for Science, Technology, and Research for the constant support during study period and preparation of the manuscript.

## Notes

### Competing Interest Statement

The authors have declared no competing interest.

